# Cultivation and genomic characterization of human gut-associated *Bdellovibrio* reveals natural predatory bacteria with specialized prey interactions

**DOI:** 10.1101/2025.11.17.688821

**Authors:** Mario Romero-Rivera, Miguel Díez-Fernández de Bobadilla, María Beltrán, Rosa del Campo, José Avendaño, Cristina Herencias

## Abstract

Predatory bacteria from the *Bdellovibrio* and Like Organisms (BALOs) group represent promising alternatives to conventional antibiotics, yet their presence in the human microbiome has remained unconfirmed through cultivation. While molecular detection methods have identified Bdellovibrionaceae DNA in human-associated environments, viable predatory bacteria had never been successfully isolated from human samples, limiting our understanding of their ecology and therapeutic potential. Here, we report the first successful isolation and characterization of viable *Bdellovibrio* strains from human fecal samples. Despite extremely low natural abundance requiring enrichment protocols for detection, two of five pooled samples yielded viable predators with characteristic lytic activity. Whole-genome sequencing of two isolates revealed >99% average nucleotide identity to the reference strain HD100 with only 26 total single nucleotide polymorphisms, indicating minimal genomic divergence between human-associated and environmental strains. Comparative genomic analysis of 163 publicly available *Bdellovibrio* genomes demonstrated that only 10.4% represented true *B. bacteriovorus sensu stricto,* highlighting substantial cryptic diversity within this genus. Pangenome analysis across 41 genomes revealed a highly conserved core genome (∼2,500-2,650 genes) contrasting with an expanding accessory genome, reflecting functional constraints of obligate predation alongside ecological adaptation. Notably, human-associated isolates exhibited narrower prey ranges, including some multidrug-resistant isolates. These findings establish that predatory bacteria are naturally associated with the human intestinal environment without acquiring novel virulence factors, support their biosafety profile for therapeutic development, and reveal prey specialization that may reflect adaptation to human microbiome ecology.

**IMPORTANCE:** This work demonstrates that bacterial predation operates as an active ecological process within the human microbiota. The successful isolation of viable *Bdellovibrio* establishes that predatory bacteria are viable microbiota members maintaining their predatory phenotype. Genomic conservation (>99% ANI to environmental strains) indicates that predation imposes stringent functional constraints superseding niche-specific adaptations. Our findings reframe microbiome understanding by revealing that predation (a top-down regulatory mechanism) operates in human-associated communities, potentially functioning as a keystone ecological process maintaining diversity through predator-prey dynamics. For microbiome research, this establishes a novel framework for host-microbe-microbe interactions where rare predators may exert disproportionate impacts on community assembly and dysbiosis pathogenesis. This work opens unprecedented opportunities for developing ecologically-informed therapeutics that harness natural predation to simultaneously combat multidrug-resistant infections while restoring microbiota homeostasis, positioning predatory bacteria as a cornerstone strategy for antimicrobial resistance management through ecological microbiota restoration.

## INTRODUCTION

The global antibiotic resistance crisis has reached alarming proportions, with one in six laboratory-confirmed bacterial infections worldwide resistant to standard treatments (1). According to the World Health Organization, antimicrobial resistance could cause up to 39 million deaths over the next 25 years, with the most vulnerable populations, including young children, the elderly, and those in low-income settings, bearing the greatest burden (1, 2). This crisis has intensified the search for novel antimicrobial strategies beyond traditional antibiotics, particularly against multidrug-resistant Gram-negative pathogens such as *Escherichia coli*, *Klebsiella pneumoniae*, *Pseudomonas aeruginosa*, and *Acinetobacter baumannii*, which constitute the most serious threats to global health (2, 3).

Among emerging alternatives, predatory bacteria belonging to the *Bdellovibrio* and Like Organisms (BALOs) group have garnered significant attention as potential “living antibiotics” (4–6). These small and motile Gram-negative bacteria exhibit a unique obligate predatory lifestyle, reproducing by invading and killing other Gram-negative bacteria through a sophisticated biphasic life cycle (4, 7–9). In natural ecosystems, including soil, freshwater, and marine environments, predatory bacteria function as keystone species that regulate bacterial populations, maintain microbial diversity, and prevent competitive dominance (mechanisms analogous to classical predator-prey dynamics that structure ecological communities) (4, 10, 11). During their attack phase, BALOs locate and attach to susceptible prey cells, penetrate the outer membrane, and enter the periplasmic space where they form a structure known as a bdelloplast. Within this protected environment, the predator consumes the host cell’s contents, replicates, and ultimately lyses the prey to release progeny that continue the predatory cycle (7, 12, 13). This obligate predatory strategy represents a fundamentally different mechanism of bacterial control than conventional antibiotics, which target specific cellular processes through chemical inhibition (9, 14, 15).

*Bdellovibrio bacteriovorus*, the most extensively studied member of this group, demonstrates remarkable therapeutic potential owing to its broad prey spectrum and ability to target clinical isolates regardless of their antibiotic resistance profiles (5, 16, 17). Unlike conventional antibiotics that target specific cellular processes, BALOs physically invade and destroy their prey, making them particularly effective against biofilms and persister cells that often evade traditional antimicrobial treatments (18, 19). Recent studies have demonstrated the safety and efficacy of *B. bacteriovorus* in *in vivo* models, showing successful reduction of pathogen loads without adverse effects on host health or immune response, including groundbreaking work in wound healing and biofilm eradication (20–23).

The known ecological distribution of BALOs extends across diverse natural environments, including soil, freshwater, marine ecosystems, and wastewater treatment facilities, where they serve as important regulators of bacterial populations (6, 8, 15, 24). Interestingly, molecular detection studies have occasionally identified BALOs in human-associated environments, including duodenal biopsies from healthy individuals and respiratory samples from cystic fibrosis patients (25, 26). These findings suggest a potential natural association between predatory bacteria and the human microbiome, although the extent and functional significance of this relationship remains largely unexplored.

The human gut microbiome represents a complex ecosystem where bacterial interactions, including predation, play crucial roles in maintaining microbial diversity and preventing pathogen overgrowth (27, 28). Theoretical frameworks suggest that predatory bacteria could function as ecological regulators within the intestinal environment, similar to their role in natural ecosystems (10, 11). However, despite compelling evidence for their presence using molecular methods, viable BALOs have never been successfully isolated from human samples, representing a significant knowledge gap in understanding their natural ecology and therapeutic potential.

The isolation and characterization of viable predatory bacteria from human sources would provide critical insights into their adaptation to host-associated environments, prey specificity patterns, and safety profiles for potential clinical applications (4, 5). Such findings would also advance our understanding of the complex predator-prey dynamics within the human microbiome and their implications for maintaining intestinal homeostasis and preventing dysbiosis-related diseases (29, 30). Furthermore, genomic characterization of human-associated BALOs could reveal adaptive features that distinguish them from environmental isolates, potentially identifying novel genetic determinants associated with host colonization, prey recognition, or survival in the intestinal environment (13, 15, 31).

Herein, we report the successful isolation of viable *B. bacteriovorus* strains from human microbiome. Through comprehensive 16S rDNA amplicon metagenomic profiling of donor microbiota, whole-genome sequencing of isolates, pangenomic analysis across 163 publicly available *Bdellovibrio* genomes, phenotypic characterization, and predation assays against clinically relevant pathogens, this work provides unprecedented insights into the natural occurrence, ecological context, and genetic features of human-associated predatory bacteria. These findings represent a significant advancement in our understanding of bacterial predation within the human microbiome and establish the foundation for future BALOs research as novel antimicrobial agents in the fight against antibiotic-resistant infections.

## RESULTS

### Detection and isolation of viable predatory bacteria associated to human microbiomes

Of the five pooled fecal samples analyzed, four (pools 1, 2, 4, and 5) yielded lytic plaques following enrichment with *P. putida* KT2440 as prey bacteria. These plaques displayed characteristic features of BALO predation: round morphology with sharp boundaries, progressive size increase over 3–5 days, and absence of bacterial colonies at their centers (**Figure 1A and Supplementary Figure 1A**). Viable predatory bacteria were quantified using the double-layer agar overlay method.

**Figure 1.**
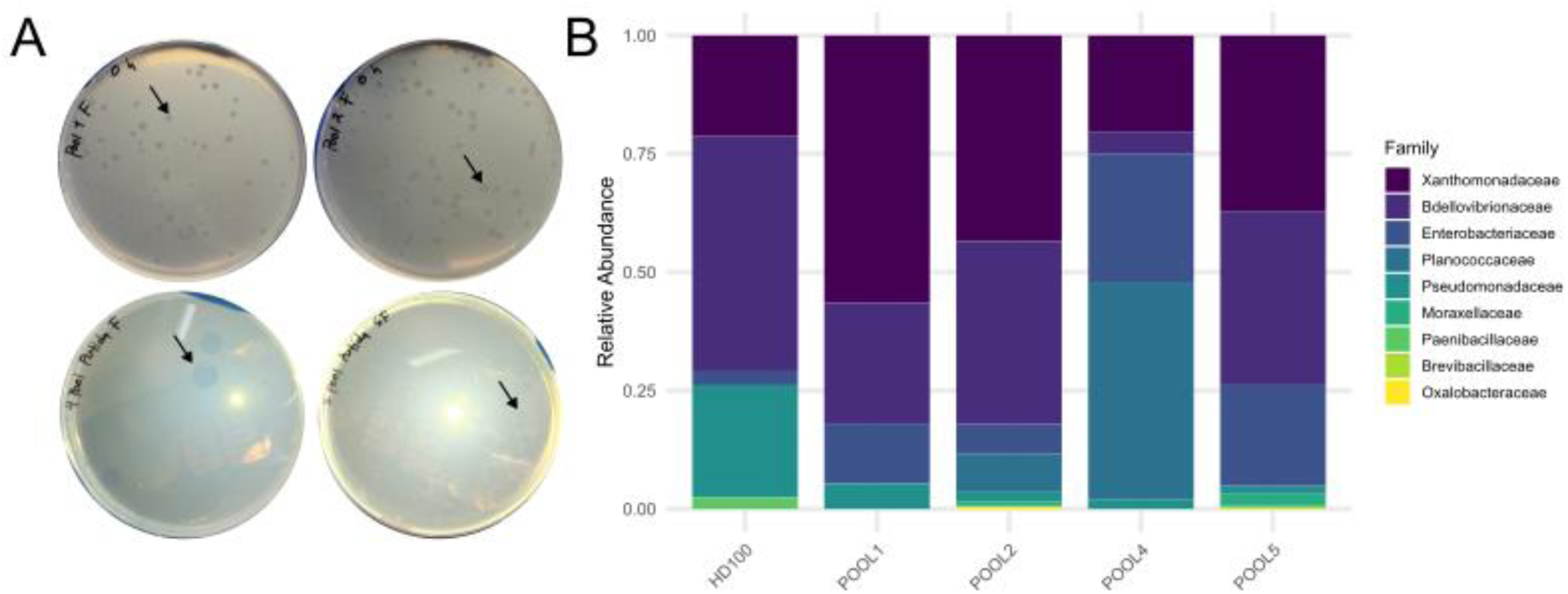
Isolation of BALOs from human fecal samples. **A)** After incubation for 72-96 h of incubation on double-layer agar culture containing P. putida, human associated predators formed circular plaques (arrows). **B)** Taxonomic composition at the family level of bacterial DNA extracted from lytic plaques formed by predatory bacteria isolated from human fecal pools 1, 2, 4, 5, and the positive control *B. bacteriovorus* HD100. Relative abundances were determined by 16S rDNA gene amplicon sequencing (V3-V4 region) analyzed using QIIME 2 and DADA2. The Bdellovibrionaceae family is highlighted, representing the predatory bacteria isolated in this study.

16S rDNA gene amplicon sequencing of DNA extracted from lytic plaques revealed variable relative abundances of Bdellovibrionaceae across plaque-derived samples: 28% in pool 1, 40% in pool 2, 5% in pool 4, 38% in pool 5, and 50% in the *B. bacteriovorus* HD100 positive control (**Figure 1B and Supplementary Figure 1B**). The remaining bacterial composition consisted primarily of prey organisms (*Pseudomonas* spp.) and commensal bacteria from the fecal matrix. Notably, despite comprehensive 16S rDNA gene sequencing of the five original fecal pools prior to enrichment, no Bdellovibrionaceae sequences were detected above the detection threshold (>10 reads per amplicon sequence variant, ASV), indicating extremely low natural abundance in the gut microbiota (**Supplementary Figure 2**). This finding is consistent with previous molecular detection studies showing transient or rare presence of BALOs in human-associated environments (25, 26). Finally, family-specific PCR amplification using primers Bd347F/Bd549R confirmed the presence of Bdellovibrionaceae DNA in Pool1 and Pool2 plaque-positive pools, yielding the expected 202 bp amplicon (25).

### Taxonomic classification of human-associated predatory bacteria

After successful enrichment in liquid medium and repeated and robust formation of lytic plaques, two of the predator isolates were selected and sequenced (from pool 1 and pool 2, hereafter BD_H1 and BD_H2, respectively). To establish taxonomic identity and phylogenetic relationships, we employed a comprehensive comparative genomic approach using the PATO pipeline (32), which integrates multiple analytical modules for bacterial genome characterization. Taxonomic assignment was performed using the PATO “classifier” function, which determines species identity by calculating average nucleotide identity (ANI) values against reference genomes in the NCBI database. The classifier assigns a reference species when ANI exceeds 95% identity. Both isolates were confirmed as *B. bacteriovorus*, with ANI values of 99.10% (BD_H1) and 99.09% (BD_H2) relative to the reference strain HD100 (**Supplementary Table 2**). These high ANI values, well above the 95% species delineation threshold, confirm conspecific status while indicating strain-level genetic variation.

The BD_H1 genome was assembled into a single circular chromosome of 3,782,514 bp with a GC content of 50.65%. The BD_H2 genome comprised three contigs with an overall GC content of 56.69%, identified through hybrid assembly using Illumina and Oxford Nanopore sequencing technologies (Contig_1, 5,354 bp; Contig_2, 3,782,519 bp; Contig_3, 5,667 bp). Detailed analysis of the smaller contigs revealed important structural and functional features. Both Contig_1 and Contig_3 harbor ribosomal RNA (rDNA) operons and transfer RNA (tRNA) genes (**Supplementary Table 3**), which are typically chromosomally encoded in bacteria.

Comparative alignment showed that Contig_1 shares 100% nucleotide identity with a specific region of the main chromosome (Contig_2) (**Supplementary Table 3**), strongly suggesting it represents a duplicated chromosomal segment or assembly artifact arising from repetitive rDNA operons rather than an independent replicon. In contrast, Contig_3 exhibited 99.75% nucleotide identity with *Stenotrophomonas maltophilia* genomic sequences (**Supplementary Table 3**), indicating a potential horizontal gene transfer (HGT) event. The presence of complete rDNA and tRNA genes on this contig, combined with its foreign origin, suggests either: (i) acquisition of a mobile genetic element carrying accessory metabolic functions during gut colonization or (ii) a genuine integrative element that has mobilized between distantly related species. Despite the presence of these additional contigs, the core genome of BD_H2 (Contig_2) exhibited typical *B. bacteriovorus* architecture and completeness metrics.

### Phylogenetic analysis of human-associated predatory bacteria

To position our human-derived isolates within the broader diversity of predatory bacteria, we retrieved 163 publicly available genomes annotated as *Bdellovibrio* spp. from the NCBI RefSeq and GenBank databases (**Supplementary Table 4**). Given the limited number of *Bdellovibrio* genomes historically available in public databases and the recent taxonomic reclassification of predatory bacteria within the phylum Bdellovibrionota (33, 34), we performed comprehensive genome-wide validation to confirm that deposited sequences truly represent authentic members of the genus *B. bacteriovorus*. To accomplish this, we employed the PATO “classifier” function, which establishes species boundaries using average nucleotide identity (ANI) calculations against rigorously validated reference genomes.

Pairwise ANI calculations between *Bdellovibrio* genomes (including our isolates BD_H1, BD_H2) revealed striking phylogenetic heterogeneity within publicly available sequences (**Supplementary Figure 3)**. Most of deposited genomes showed low ANI values (<90%), indicating they represent distinct species or even genera within the Bdellovibrionaceae family. This finding highlights the substantial taxonomic diversity that remains poorly characterized within this predatory bacterial group and underscores the need for systematic genomic-based reclassification of historical isolates.

Applying the widely accepted >94% ANI threshold for bacterial species delineation (35), we identified only 17 genomes (10.4% of total) that qualified as *B. bacteriovorus sensu stricto*. To examine fine-scale genomic variation within this species complex, we performed a single nucleotide polymorphism (SNP) analysis across the core genome and calculated SNP density normalized per megabase (SNPs/Mb) for all pairwise comparisons (**Figure 2A and Supplementary Table 5**). SNP-based heatmap analysis revealed distinct patterns of genomic divergence within the *B. bacteriovorus* clade. BD_H1 and BD_H2 exhibited remarkably low SNP densities when compared to each other and to the reference strain HD100 (**Supplementary Table 2**). These extremely low SNP densities indicate a recent common ancestry and minimal evolutionary divergence between human-derived isolates and well-characterized environmental strains.

**Figure 2.**
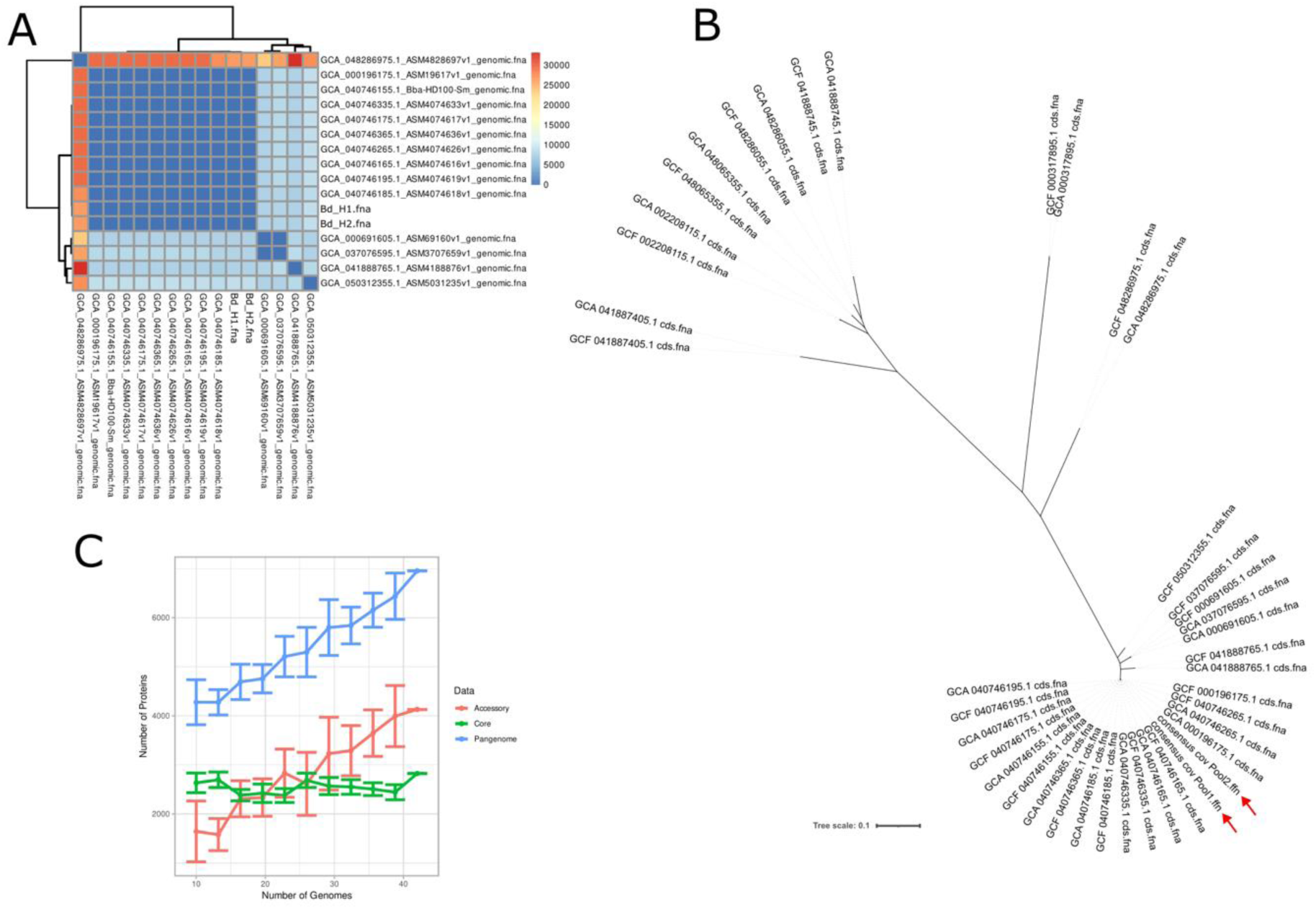
Genomic characterization and comparative analyses of 41 *Bdellovibrio* strains. **A)** Heatmap showing pairwise core genome SNP counts per megabase calculated across 17 genomes with an ANI ≥94%. The accession numbers are listed on the axes. **B)** Maximum likelihood phylogenetic tree based on core genome SNPs (n=41 genomes). The tree topology recapitulates the ANI-based clustering, with distinct branching patterns separating *B. bacteriovorus* sensu stricto (short internal branches, indicating recent divergence) from the more distantly related 90–94% ANI group (longer branches). The tree demonstrates that both phylogenetic (SNP-based) and genomic similarity (ANI-based) metrics converge on the same population structure, validating the taxonomic classification. Red arrows point to the human-associated *Bdellovibrio*. **C)** Pangenome composition and scaling dynamics. Plot showing the accumulation of core genes (green line), accessory genes (red line), and total pangenome size (blue line) as a function of sequenced genomes (n=41). Error bars represent 95% confidence intervals from 100 rarefaction replicates.

To further resolve the relationships within the *B. bacteriovorus* sensu stricto and its closest relatives, we selected the 17 genomes with ANI ≥95% and an additional 24 genomes with ANI of 90–94%, for a total of 41 high-quality genomes (**Supplementary Figure 3**). We then performed a core genome analysis and constructed a matrix of pairwise average nucleotide identity (ANI) values, alongside single nucleotide polymorphism (SNP)-based phylogenetic analysis of the core annotated genome (**Supplementary Figure 4 and Supplementary Figure 5**).

Complementing the ANI-based clustering, the SNP-based maximum likelihood phylogenetic tree of the core genome (**Figure 2B**) provides additional resolution of relationships among these 41 genomes. The human-derived isolates clustered together with HD100 and closely related reference strains, supporting both their classification and their close evolutionary relationship, while other clades showed longer branches, indicative of additional divergence.

We then analyzed the pangenome composition and evolution across the 41 *Bdellovibrio* genomes, stratified by core genome, accessory genome, and total pangenomes (**Figure 2C**). The core genome (green line), representing genes present in ≥95% of the genomes, remained remarkably stable and nearly constant across all 41 analyzed strains, plateauing at approximately 2,500–2,650 orthologous gene families. This genomic invariance, even across geographically and ecologically diverse isolates (including human-derived, environmental aquatic, soil, and wastewater strains), indicates strong functional constraints on the predatory lifestyle. The obligate predatory strategy, which requires specialized systems for prey sensing, attachment, invasion, and intracellular lysis, appears to impose a well-defined minimal genomic blueprint that cannot be substantially altered without compromising fitness. In contrast, the accessory genome (red line) exhibited substantial variation and consistent expansion as more genomes were sequenced, reaching approximately 4,100–4,200 gene families by the 41 genomes. The steep trajectory of the red curve indicates that each newly added genome contributes an average of ∼60–100 novel genes not found in the previously sequenced strains. Finally, the total pangenome (blue line) demonstrated logarithmic growth, increasing from ∼4,000 genes in the first 10 genomes to ∼6,500–7,000 genes at n=41, approaching saturation but not yet fully converging. This incomplete saturation suggests that additional *Bdellovibrio* diversity remains undescribed, consistent with our finding that only 10.4% of deposited *Bdellovibrio*-labeled genomes are truly *B. bacteriovorus* sensu stricto. The continued accumulation of novel accessory genes underscores the substantial hidden diversity within this genus and supports the need for systematic taxonomy-guided exploration and characterization of predatory bacterial isolates from undersampled ecological niches.

The remaining 122 genomes (74.8%) displayed ANI values below 90%, confirming their assignment to divergent lineages within the broader Bdellovibrionaceae family. This distribution reveals that current NCBI annotations significantly oversimplify the taxonomic complexity of predatory bacteria, with many genomes labeled as “*Bdellovibrio* sp.” actually representing phylogenetically distant taxa. Our analysis demonstrated that the genus *Bdellovibrio* encompasses far greater genomic diversity than previously recognized, with multiple cryptic species awaiting formal description.

### Predatory activity and prey specificity of human-associated BALOs

To assess prey range and specificity, both human-derived BDPool1 and BDPool2 predators and the reference strain *B. bacteriovorus* HD100 were challenged against a panel of clinically relevant prey organisms in three to five independent replicate predation assays (**Table 1 and Supplementary Table 6**).

**Table 1.** Predatory capability of *B. bacteriovorus* HD100 and BALOs isolates from BD_H1 and BD_H2. Raw data of PFU are compiled in Supplementary Table 6. +; for predator-prey combinations with plaque formation in at least one replicate; +++, for predator-prey combinations with >10 plaque formations in more than one replicate.

| Prey Strain | HD100 | BD_H1 | BD_H2 |
| --- | --- | --- | --- |
| *E. coli* ATCC 25922 | + | - | - |
| *K. pneumoniae* CR | + | - | - |
| *K. pneumoniae* CS | + | - | - |
| *P. aeruginosa* 14 | + | - | + |
| *P. aeruginosa* 14M | + | + | + |
| *P. aeruginosa* 28 | + | + | - |
| *P. aeruginosa* 28M | + | - | - |
| *P. aeruginosa* 40 | + | + | - |
| *P. aeruginosa* 40M | - | + | - |
| *P. aeruginosa* 7 | - | - | - |
| *P. aeruginosa* 7M | + | + | - |
| *P. aeruginosa* PAO1 | - | + | - |
| *P. aeruginosa* PAO1 M | + | + | - |
| *P. putida* KT2440 | +++ | + | + |

Predation assays revealed marked strain-specific prey preferences among the three predators tested. The reference strain HD100 demonstrated the broadest prey spectrum, efficiently preying on *P. aeruginosa*, *P. putida*, *E. coli*, and select *K. pneumoniae* isolates. In contrast, BD_H1 and BD_H2 predators exhibited narrower prey ranges, with their activity predominantly restricted to Pseudomonas spp. Neither BD_H1 nor BD_H2 predators showed detectable lytic activity against *K. pneumoniae* or *E. coli* isolates tested.

Variability between replicate experiments was observed for certain predator-prey combinations, likely reflecting the dynamic nature of predatory interactions and potential heterogeneity in prey surface structures. This variability underscores the importance of standardized assay conditions and replicate testing when evaluating predatory phenotypes (7, 36).

## DISCUSSION

The successful isolation of viable *B. bacteriovorus* from human fecal samples represents a significant milestone in predatory bacterial research, bridging the gap between molecular detection studies and actual cultivation of these organisms from human sources. While previous work by Iebba and colleagues demonstrated through PCR and quantitative PCR that *Bdellovibrio* DNA was present in human intestinal biopsies with significantly higher prevalence in healthy controls (100%) compared to patients with inflammatory bowel disease and celiac disease (11-43%) (25), and more recently metagenomic profiling identified Bdellovibrionaceae sequences in complex respiratory microbiomes such as cystic fibrosis lung tissue (26), our work now establishes that predators are not merely transient DNA signatures but viable, metabolically active members of the human microbiota capable of maintaining their predatory activity under appropriate conditions (**Table 1**).

Our findings demonstrate that despite extremely low natural abundance in the intestinal environment (where Bdellovibrionaceae were undetectable in the original samples prior to enrichment, as shown in **Supplementary Figure 2**), viable predatory bacteria are associated with the human microbiome and may play ecological roles in microbial community regulation. This finding aligns with recent advances in culturomics showing that approximately 70-80% of the human gut microbiota can be cultured using diverse enrichment approaches, enabling recovery of numerically rare taxa that exert disproportionate functional impacts on community structure and ecosystem function (37, 38). The necessity of enrichment protocols reflects that predatory bacteria, like many ecological specialists, occupy specific microniches and may exist in density-dependent equilibrium with prey populations through frequency-dependent predator-prey dynamics rather than constituting dominant community members (39, 40).

The isolation of Bd_H1 and Bd_H2 with genomic signatures nearly identical to well-characterized environmental strains (ANI >99% to HD100, with minimal SNP densities of 26 total polymorphisms across both isolates) is particularly significant for biosafety considerations, as it indicates that human-associated predators have not acquired novel virulence factors or substantially diverged from laboratory, reference, or environmental strains. Our comparative genomic analysis of 163 *Bdellovibrio*-labeled sequences revealed striking taxonomic heterogeneity within public databases, with only 10.4% qualifying as true *B. bacteriovorus* sensu stricto based on the 94% ANI species delineation threshold (**Supplementary Figure 3**). This finding underscores the substantial cryptic diversity within predatory bacteria that remains poorly characterized and formally undescribed, highlighting the importance of rigorous genome-based taxonomy and systematic exploration of predatory isolates from diverse ecological niches. The pangenome analysis across 41 genomes demonstrated a paradoxical pattern: a highly conserved core genome (∼2,500-2,650 genes) reflecting stringent functional constraints imposed by the obligate predatory strategy, combined with substantial expansion of the accessory genome. This pattern suggests that while the fundamental predatory machinery is tightly constrained, individual strains acquire diverse metabolic and regulatory capabilities, potentially reflecting adaptation to specific ecological niches or prey populations.

The observed strain-specific prey preferences constitute perhaps the most functionally significant finding, with Bd_H1 and Bd_H2 isolates demonstrating markedly narrower prey ranges restricted primarily to *Pseudomonas* spp., whereas reference strain HD100 efficiently predated *Pseudomonas*, *Escherichia*, and *Klebsiella* spp.. This specialization may reflect ecological adaptation to the human gut environment or selection during laboratory enrichment of *P. putida* prey and suggests that prey range determination is multifactorial and potentially influenced by lipopolysaccharide structure, particularly lipid A composition and modifications. Critically, variable predation efficiencies against isogenic pairs of clinical isolates differed in documented LPS modifications(41, 42). Particularly *P. aeruginosa* strains with murepavadin resistance conferred by lipid A acylation changes (42), and *Klebsiella pneumoniae* isolates from urinary tract infections, and *K. pneumoniae* strains with colistin resistance due to 4-amino-arabinose addition to lipid A phosphate groups (41), suggesting that antimicrobial resistance-associated surface modifications can mechanistically influence predator attachment efficiency and invasion kinetics. These findings suggest that bacteria acquiring antibiotic resistance through outer membrane remodeling paradoxically create or abolish predator epitopes, establishing an ecological link between antibiotic resistance evolution and predation susceptibility with potentially significant implications for sequential evolution of multiple defense mechanisms. The therapeutic implications of narrow prey specificity present potential advantage over broad-spectrum activity for targeted applications. This approach could be directed against specific ESKAPE pathogens, particularly *P. aeruginosa*, a leading cause of healthcare-associated infections and the dominant pathogen in cystic fibrosis lung disease (43).

Notably, genomic analysis identified a contig of 99.75% nucleotide identity to *S. maltophilia* genomic sequences in isolate BD_H2, containing complete rDNA and tRNA operons. While this finding could represent contamination or assembly artifacts, an alternative hypothesis warrants consideration, as these sequences may be derived from prey nucleotides incorporated during *Bdellovibrio* DNA replication within the bdelloplasts. Classic radiolabeling experiments using (^14^C)uracil-labeled *E. coli* demonstrated that approximately 50% of prey RNA radioactivity is incorporated into *Bdellovibrio* RNA with identical specific activity, and prey DNA contributed substantially to predator DNA synthesis, with 73% of ^3^H-thymidine from labeled prey incorporated into *Bdellovibrio* DNA (44, 45). Given that BD_H2 was enriched through serial predation on Pseudomonas spp., the detected *S. maltophilia* sequences (if representing salvaged prey nucleotides) would suggest natural predation on this organism in the original human microbiota prior to isolation. This interpretation gains ecological significance when contextualized within documented presence of *S. maltophilia* in human fecal microbiota: 10.9% carriage in diarrheal patients and 33% in hematologic malignancies (46, 47), recent metagenomic detection in colorectal cancer intestinal samples, and increased respiratory colonization in cystic fibrosis (3% to 15%).

This study provides critical insights into the natural ecology of predatory bacteria in the human microbiome and establishes a foundation for evaluating their therapeutic potential. Critically, the isolation of human-derived predators represents a paradigm shift in predatory bacteria research. To date, essentially all laboratory and therapeutic development studies investigating *B. bacteriovorus* have employed isolates derived from environmental sources (primarily soil and freshwater ecosystems) or reference strains maintained through decades of laboratory cultivation (21, 23, 48). These environmentally-sourced or laboratory-adapted strains, while demonstrating impressive antimicrobial activity *in vitro* and efficacy in animal infection models, were never naturally selected for persistence or predatory activity within human-associated microbiota. The isolation of BD_H1 and BD_H2 directly from human fecal samples, coupled with their demonstration of viability, functional predatory capacity, genomic integrity (>99% ANI to reference strains), and activity against clinically-relevant multidrug-resistant pathogens, opens an unprecedented opportunity: the possibility of developing *Bdellovibrio*-based therapeutics using naturally human-associated strains that may be inherently optimized for survival, predatory activity, and immunological compatibility within human biological systems (5, 49). The transition from environmental isolates to human-derived predatory bacteria as therapeutic agents could substantially enhance clinical translatability while simultaneously advancing our understanding of predatory bacteria ecology, prey-predator coevolution, and the role of top-down regulation in human microbiome homeostasis, ultimately positioning predatory bacteria as a revolutionary biocontrol strategy for the global antimicrobial resistance crisis. Beyond direct antimicrobial activity, human-derived *Bdellovibrio* may function as ecological regulators within the microbiota, potentially restoring predator-prey balance in dysbiotic states and contributing to microbiome-mediated therapeutic strategies.

## MATERIALS AND METHODS

### Ethical consideration and human fecal samples

This study was conducted in accordance with the ethical standards of Hospital Universitario Ramón y Cajal. Fifty anonymized, non-diarrheal stool samples previously submitted to the Microbiology Department for routine parasitological screening were collected without identifiable patient information. Samples were pooled into five groups of ten individual specimens each to facilitate BALO enrichment and detection.

### Bacterial strains and culture conditions

*B. bacteriovorus* HD100 (ATCC 15356) was used as control predator strain and *Pseudomonas putida* KT2440 as the reference prey (50). Additional prey strains included *P. aeruginosa* PAO1 (ATCC 47085) and *Escherichia coli (*ATCC 25922), and clinical isolates of *P. aeruginosa* obtained from cystic fibrosis patients (42), and *Klebsiella pneumoniae* isolates from urinary tract infections (41). Clinical strains were selected based on their antibiotic resistance profiles and previously documented lipopolysaccharide modifications. The strains used in this study are listed in **Supplementary Table 1**.

The prey strains were cultured from glycerol stock in blood agar (CAN-I-BD) medium at 37°C and were cultivated in liquid with shaking at 180 rpm in LB medium. All predation co-cultures were performed in diluted nutrient broth (DNB, 1:10 dilution of standard nutrient broth) supplemented with 2 mM CaCl₂ and 1 mM MgSO₄ (51).

### BALO enrichment from fecal samples

BALO enrichment was performed following an adaptation of established protocols (25, 51). Ten grams of pooled fecal material were suspended in 50 mL of sterile saline solution (0.9% NaCl) and vortexed vigorously for 5 min to homogenize. To this suspension, 5 mL of DNB containing 10^8^ CFU/mL of *P. putida* KT2440 was added as prey. The enrichment culture was incubated at 30°C with shaking at 150 rpm for 48 h to allow the growth of predatory bacteria.

Following incubation, cultures were filtered through 0.45 µm cellulose acetate membrane filters (Millipore) to remove the prey bacteria and retain potential predatory bacteria in the filtrate. One milliliter of filtrate was transferred to tubes containing 500 µL of each potential prey strain adjusted to 10^8^ CFU/mL and co-incubated for 24 h at 30°C and shaking 150 rpm.

### Double-layer agar overlay assay for BALO enumeration

BALO viability and enumeration were determined using the double-layer agar overlay method (52). Briefly, 1.5 mL of co-culture was added to 4 ml of melted DNB soft agar (0.7% w/v), maintained at 50°C, supplemented with 2 mM CaCl₂ and 1 mM MgSO₄. The mixture was gently vortexed and immediately poured onto plates containing DNB agar (1.5 % w/v). Plates were allowed to solidify at room temperature for 30 min before incubation at 30°C for 48–72 h. Clear lytic plaques as indicative of predatory activity were counted and expressed as plaque-forming units per milliliter (PFU/mL).

### DNA extraction and molecular confirmation of BALOs

Genomic DNA was extracted from fecal pools and purified lytic plaques using the QIAamp DNA Stool Mini Kit (Qiagen, Germany) following the manufacturer’s protocol. For fecal samples, we used an aliquot of 3 mL, centrifuged and proceeded with the extraction protocol. For lytic plaque samples (Pool 1, 2, 4, 5), plaques were excised from soft agar, resuspended in 500 µL of sterile phosphate-buffered saline, and centrifuged at 10,000 × g for 10 min to pellet cells before proceeding with the extraction protocol. DNA concentration and purity were assessed using a NanoDrop spectrophotometer (Thermo Fisher Scientific), and DNA integrity was evaluated by agarose gel electrophoresis. Extracted DNA was stored at −20°C until further analysis.

To confirm the presence of predatory bacteria, PCR amplification targeting the Bdellovibrionaceae family was performed using specific primers Bd347F (5’-ATAAGGGATGACGACGACGGAGG-3’) and Bd549R (5’-GCTAG GATCCCTCGTCTTACC-3’), following the conditions previously established (25).

### 16S rDNA gene amplicon sequencing

To characterize the bacterial composition of fecal pools and lytic plaques, 16S rDNA gene amplicon sequencing targeting the V3-V4 hypervariable region was performed. PCR amplification was conducted using universal primers 341F (5’-CCTACGGGNGGCWGCAG-3’) and 805R (5’-GACTACHVGGGTATCTAATCC-3’) with Illumina adapter sequences. Amplicons were sequenced on an Illumina MiSeq platform using the MiSeq Reagent Kit v3 (2 × 300 bp) for 600 cycles.

Raw sequencing reads were processed using the QIIME 2 bioinformatics platform (52). Quality filtering, denoising, and ASV identification were performed using the DADA2 (53). Forward and reverse reads were trimmed to remove primers and low-quality bases, and reads were merged with a minimum overlap of 12 bp. Chimeric sequences were identified and removed using the consensus method in DADA2. Taxonomic classification of ASVs was performed using the SILVA 138 database (54) with the classify-sklearn naive Bayes classifier. ASVs with total abundances below 10 reads across all samples were removed to control for spurious sequences and sequencing errors.

### Whole-genome sequencing and assembly

Due to the low abundance of predatory bacteria in co-cultures with prey, DNA enrichment was necessary prior to whole-genome sequencing. Isolated BALOs from pools 1 and 2 were subjected to sequential enrichment by co-culturing with increasing volumes of *P. putida* prey. Specifically, predators were incubated with 1 mL of prey (10⁸ CFU/mL) in 8 mL HEPES buffer (25 mM, pH 7.8) at 30°C with shaking at 170 rpm for 72 h. Cultures were centrifuged at 15,000 × g for 5 min at room temperature, supernatants were discarded, and pellets were resuspended in double the previous prey volume with fresh HEPES buffer. This process was repeated four times until sufficient biomass was obtained, as evidenced by visible pellet formation.

Total genomic DNA was extracted using the QIAamp DNA Mini Kit (Qiagen) and quantified using the Qubit dsDNA HS Assay Kit (Thermo Fisher Scientific). For short-read sequencing, DNA libraries were prepared using the Nextera XT DNA Library Preparation Kit (Illumina) and sequenced on an Illumina MiSeq platform with the MiSeq Reagent Kit v2 Nano (2 × 150 bp) for 300 cycles, generating 250,000–500,000 reads per sample. Genomic DNA were sequenced with Oxford Nanopore technology at Plasmidsaurus (https://plasmidsaurus.com/, Eugene, USA) using the V10 chemistry library prep kit with the R10.4.1 flow cells. Sequences were annotated using BAKTA (55).

### Genomic analysis, Taxonomic typing and core genome

The 163 deposited genomes were initially annotated using Bakta 1.7.0 (55). Subsequently, core and accessory genomes were defined using the PATO tool (32). This allowed for an analysis of the population structure, the annotation of adaptive features, and the creation of gene networks. Specifically, we utilized the PATO functions “core_plots” to determine the size of the pangenome, core, and accessory genome, and “similarity_tree” to generate pseudo-phylogenetic trees.

We assigned the species to each isolated using the function “classifier” of PATO(32). “Classifier” assigns each query genome to the closest reference genome from the NCBI (https://www.ncbi.nlm.nih.gov/taxonomy) by calculating the ANI of each genome to the reference ones. It assigns a reference species if the ANI is over 95% of identity. The heatmap from **Supplementary Figure 3** that included 163 sequenced genomes of *Bdellovibrio* from the NCBI was created using the function “similarity_tree” and the values from the function “mash” from PATO. Clades were defined based on minimum within-node pairwise ANI scores using the unsupervised clustering MClust (56).

## Data availability

Assembled genome sequences for isolates BD_H1 and BD_H2 have been submitted to the EBI database under the project accession number PRJEB102169.

## ACKNOWLEDGMENTS

Work in the Metagenomics lab is supported by PRECICOLON P2022/BMD-7212 from Comunidad de Madrid, METOXISAN project from Fundación Mutua Madrileña, END-RAM PLEC2024011123 project by Agencia Estatal de Investigación and MePRAM PMP22/00092 funded by Instituto de Salud Carlos III. Work of MRR was supported by Fundación Carolina (C2022). During implementation of this study, MDFdB was supported by the Instituto de Salud Carlos III (pFIS F19/00366). CH is supported by the European Union (ERC, HorizonGT, 101077809), PI23/01945 by the Carlos III Health Institute (ISCIII), and FERP-2024-182 (2024/0357) funded by Fundación Rodríguez Pascual.

